# A *Leptosphaeria maculans* set of isolates characterised on all available differentials and used as control to identify virulence frequencies in a current French population

**DOI:** 10.1101/2020.01.09.900167

**Authors:** L. Bousset, M. Ermel, R. Delourme

## Abstract

The characterization of virulence frequencies has to be regularly updated to identify which genes are currently efficient and use this information to advise gene deployment by choosing varieties depending on the current composition of local pathogen population. In *L. maculans* on *Brassica napus*, because different genes were characterized by different teams, because new interactions are continuously identified and seed of differentials are difficult to obtain, we today still lack isolates characterized on all current resistance genes. On the one hand, we assembled a set of 12 isolates characterized on 13 of the 17 described resistance genes, having clearly compatible and clearly incompatible isolates for each interaction. This set can be used to characterize the *L. maculans* – *B. napus* interaction at cotyledon stage. Expanding the set of isolates with clearly virulent ones allowed us to detect inconsistent behaviour or intermediate (avirulent) phenotypes. On the other hand, we used this set of isolates as controls to identify virulence frequencies in a current French *L. maculans* population sampled in 2018 at Le Rheu. We provide the current status for 13 avirulence frequencies, including *LepR1*, *LepR2* and *LepR3* available in near isogenic lines of spring canola but not yet documented in France. Avirulence frequencies on the genes *Rlm1*, *Rlm2*, *Rlm3*, *Rlm4*, *Rlm7*, *Rlm9* and *LepR3* were low, indicating the lack of efficacy of these genes against the current population. In the opposite, all or most of isolates were avirulent for the genes *Rlm5*, *Rlm6*, *Rlm10*, *Rlm11*, *LepR1* and *LepR2*. An optimistic point of view could conclude that there are ample resources for oilseed rape breeding. However, as compared to previous studies, so far all the resistance genes used on significant acreage without additional management practices have lost efficacy and only avirulences corresponding to resistance genes not deployed in France retain efficacy. While the call to wisely manage the available host resistance genes is not recent, it is still relevant. Adding, management practices to the deployment of resistance genes in order to reduce inoculum carry-over from one growing season to the next and to lower population sizes is key to maintain their efficacy over time.

## Introduction

In crops, breeding varieties for increased disease resistance is a major means to control epidemics, even if focusing on resistance to one disease could inadvertently increase susceptibility to another (Arraiano & Brown, 2017). However, the deployment of different resistance genes through space and time drives the genetic composition of the pathogen population (Hovmøller et al., 1997; Papaïx et al., 2011). Because epidemics of successive cropping seasons are not independent, the adaptation of pathogen populations to host resistances proceeds at the scale of a network of fields on which the selection pressures are not homogeneous, during a succession of cropping seasons (Bousset & Chèvre, 2013). The call to wisely manage the available host resistance genes is not recent, but still relevant. It is today understood that passively maintaining resistance efficacy is unlikely. The loss of efficacy of major genes depends on their deployment, and theoretical knowledge allow to link resistance deployment and pathogen population adaptation. Previous studies indicate that resistance genes in varieties exert directional selection for isolate-host compatibility (Hovmøller et al., 1993, Bousset et al. 2018). Predictable changes in pathotype frequencies were observed following commercial use of genes, or in field experiments (Hovmøller et al., 1993, Brun et al., 2010, Daverdin et al., 2012). This opens the perspective of actively maintaining resistance efficacy by choosing varietal deployment depending on the current composition of local pathogen population. Thus, the characterization of virulence frequencies has to be regularly updated to identify which genes are currently efficient, and use this information to advise gene deployment (Van de Wouw et al. 2014).

Breeding for resistance to fungal pathogens contributed to yield increase in oilseed rape (Potter et al., 2016) and is the main means to control phoma stem canker. *Leptosphaeria maculans* is the cause of stem canker on oilseed and vegetable brassicas, especially *B. napus*, *B. juncea* and *B. oleracea* (Mendes-Pereira et al., 2003). Epidemics are initiated in autumn, mainly by wind dispersed ascospores produced following sexual reproduction on stubble (Lô-Pelzer et al., 2009). Leaf spots are observed from autumn to early spring. Stem cankers develop from spring to summer, up to the time of harvest, due to the systemic growth of the fungal hyphae from leaf spots to the leaf petiole through vessels, and subsequently to the stem base (Travadon *et al*., 2009). In *B. napus* there are more than 17 qualitative resistance genes conferring resistance to *L. maculans* (see Delourme et al., 2006; Rimmer, 2006; Elliott et al. 2015; Raman et al., 2013 for reviews and Larkan et al. 2019). Their presence in host varieties can be detected using sets of *L. maculans* isolates to inoculate seedlings under controlled conditions. Phoma stem canker populations are diversified worldwide, with contrasted histories regarding the use of plant resistance genes (Rouxel et al., 2003a, Marcroft et al., 2012, Zhang et al., 2016). Populations of *L. maculans* collected from hosts with different resistance gene combinations harbor different virulence allele frequencies (Van de Wouw et al., 2014, 2017). When resistance genes have been deployed in oilseed rape varieties, phoma stem canker populations have repeatedly become adapted over a few years (Balesdent et al. 2006; Liban et al., 2016, Van de Wouw et al. 2017). Documented examples are adaptation to the *B. napus* gene *Rlm1* in Europe (Rouxel et al., 2003b); the genes contained in the variety Surpass 400 in Australia (Li et al. 2003; Van de Wouw et al., 2014, 2017); *Rlm7* in Europe (Leflon, 2013; Winter & Koopmann 2016); and *Rlm3* in Canada (Zhang et al., 2016). Thus, *L. maculans* has repeatedly evolved virulence against nearly all major resistance genes released in oilseed rape so far, sometimes with almost complete yield loss.

With so many genes, large scale survey on many populations becomes impractical. One can hope that easy to use sets of molecular markers can be developed. Nevertheless, phenotype cannot always be deduced from the genotype, because of interference between avirulence genes. For example, avirulence at the Avr4-7 loci masks avirulence at the Avr3 locus (Plissonneau et al. 2016) and at the Avr5-9 locus (Ghanbarnia et al. 2018); loss of avirulence at the Avr1 locus masks avirulence at the AvrLepR3 locus (Larkan et al. 2013). The need to phenotype populations remains, to identify additional cases in which such interferences might occur, or new alleles. Because different genes have been characterized by different teams, because new interactions are continuously identified (Larkan et al. 2019) and seed of differentials are difficult to obtain, we today still lack isolates characterized on all current sets of genes. Choosing isolates with already published information and expanding the set to encompass clear contrasted phenotypes, our first aim was to expand the characterization of a set of isolates on as many avirulence genes as possible, in order to use them as controls when testing a population of isolates.

When no previous data on gene efficacy are available, gaining knowledge on at least a population might provide useful information to optimize the breeding effort. In France, populations have been surveyed following the release and subsequent loss of efficacy of *Rlm1* (Rouxel et al. 2003b; Balesdent et al. 2005, 2006). At that time virulence frequencies towards *Rlm6*, *Rlm7* and *Rlm10* were below 1%. Following the release of *Rlm7* in commercial varieties, the frequency of compatible isolates has increased up to locally 50% (Balesdent et al. 2015; 2019; Leflon 2013). In Canada and Australia, isolates compatible on *Rlm6* have been detected in *L. maculans* populations at frequencies of 6% and up to 70% depending on the year (Liban et al. 2016; Van de Wouw et al. 2017), and it remains to be tested if *Rlm6* and *Rlm10* retain efficacy in France. Specific studies have documented frequencies towards some genes, such as *Rlm11* with frequencies of compatible isolates at Le Rheu as low as 3 out of 22 isolates in 2000-2001 and none of the 32 isolates virulent in 2010-2011 (Balesdent et al. 2013). Despite the development of near isogenic lines for *LepR1*, *LepR2* and *LepR3* (Larkan et al. 2016) we still lack information about their efficacy in France. Using as many genes as possible, our second aim was to characterize their efficacy in a current French *L. maculans* population, so that wise use of the currently efficient ones could be proposed.

## Materials and methods

### Fungal population to identify avirulence gene frequencies

A succession of trays of 10 days old seedlings of the susceptible variety Westar (no resistance genes) were exposed to the airborne inoculum at Le Rheu (Brittany, France) from October 2017 until January 2018. On 16 January 2018 (no leaf spots had been observed before), *L. maculans* leaf spot population was sampled by collecting diseased cotyledons. From each cotyledon, one typical lesion was excised and placed on wet absorbent paper in a Petri dish. After allowing the lesion to sporulate for 24 h at room temperature, 30 single pycnidial isolates per plot were produced (one isolate per leaf spot). Spores oozing from single pycnidia were collected with a sterile needle under a magnifying lens and transferred onto malt-agar.

Inoculum, consisting of suspensions of 10^7^ pycnidiospores per mL, was obtained for each isolate. Specifically, V8 agar (V8 juice 160 ml.l^−1^, agar 20 g.l^−1^) was autoclaved for 20 min at 120°C. After cooling, streptomycin concentrated solution 10% w/v in water was added to a final concentration of 0.1 g.l^−1^. For each isolate, agar plugs were transferred to V8 agar and grown for 10 days under near UV light. Pycnidiospores were dislodged from plates in sterile distilled water, filtered through muslin cloth. Spore concentrations were standardized to 107 spores per ml after counting aliquots with a Malassez cell under a magnifying lens, aliquoted and stored at −20°C until tested for virulence.

### Plants and inoculation

A set of 17 oilseed rape lines or varieties with known resistance genes were used (Table 1). After pregermination on wet filter paper, seeds of each variety were transplanted in trays with a 1:1:1 mix of sand, peat and compost and grown in climate chamber under a 16-h photoperiod for 10-12 days, at temperatures of 18°C night / 20°C day. A 10 µL drop of 10^7^ conidia per ml suspension was placed on each lobe of prick-wounded cotyledons (4 inoculation sites per plant) for six plants of each variety, in two independent trays. A plastic cover was placed over the inoculated plants to create a 100% relative humidity (RH) atmosphere for 24h in the dark.

**Table 1.** Presence of the resistance alleles in the differential lines used.

| Lines | Genes | Reference |
| --- | --- | --- |
| MT29 | *RLm1 + Rlm9* | Delourme et al. 2008 |
| Bristol | *Rlm2 + Rlm9* | Balesdent et al. 2005 |
| Grizzly | *Rlm3 + Rlm2* | Jestin et al. 2015 |
| Falcon | *Rlm4* | Rouxel et al. 2003 |
| Darmor | *Rlm9* | Delourme et al. 2004 |
| 99.150.2.1 | *Rlm5* | Stachowiak et al. 2006 |
| Eurol-*Rlm6*<sup>a</sup> | *RLm6 + Rlm2 + Rlm3* | Brun et al. 2010 |
| Falcon-*Rlm6*<sup>a</sup> | *RLm6 + Rlm4* | Balesdent et al. 2002 |
| 01.23.2.1 | *RLm7* | Delourme et al. 2004 |
| Eurol-*Rlm10*<sup>b</sup> | *Rlm10 + Rlm2? + Rlm3?*<sup>c</sup> | Petit-Houdenot et al. 2019 |
| Westar-*Rlm10* | *Rlm10* |  |
| Darmor-*Rlm11* | *Rlm11*<sup>d</sup> | Balesdent et al. 2013 |
| Topas-*LepR1* | *LepR1* | Larkan et al. 2016 |
| Topas-*LepR2* | *LepR2* | Larkan et al. 2016 |
| Topas-*LepR3* | *LepR3* | Larkan et al. 2016 |
| Surpass 400 | *LepR2 + LepR3*<sup>e</sup> | Yu et al. 2008 |
| Napoli | *LepR2*<sup>f</sup> | Terres-Inovia 2019 |

<sup>a</sup> Eurol-*Rlm6* and Falcon-*Rlm6* were also named EurolMX and FalconMX, respectively.
<sup>b</sup> Eurol-*Rlm10* was also named FuEurol74.
<sup>c</sup> *Rlm2* and *Rlm3* genes are present in the Eurol progenitor of Eurol-*Rlm10*. Because we currently lack pathogen isolates both virulent on *Rlm10* and avirulent on *Rlm2* and *Rlm3*, the presence of these two genes in Eurol-*Rlm10* has not yet been tested for.
<sup>d</sup> *Rlm9* is present in the Darmor progenitor of Darmor *Rlm11* and its absence in the Darmor *Rlm11* line is confirmed in this study (see discussion).
<sup>e</sup> the gene content of Surpass 400 is much debated (see discussion).
<sup>f</sup> Napoli was published having *RlmS* (Terres-Inovia 2019; see results)

### Fungal reference isolates

At least two reference isolates with known phenotypes were included as controls in each test, so that both compatible (virulent) and incompatible (avirulent) reactions were generated and the expression of interactions was confirmed. Some prior information on isolates’ genotypes was known from previous publications (Table 2), and our study allowed expanding the information for genes not previously determined (Table 3).

**Table 2.** Origin and source of prior information for the 12 fungal reference isolates.

| Isolate | Origin | Year | Other name | Source of prior information |
| --- | --- | --- | --- | --- |
| 00-100 | Canada | 2000 |  | a; b; c |
| JN2 | France | Cross | v23.1.2 | d; e; f; g; h |
| JN3 | France | Cross | v23.1.3 | b; c; d; f; g; h;i |
| IBCN14 | Australia | 1988 | MD2 | f; i; j; k; l |
| IBCN16 | Australia | 1988 | 10 | j; m |
| IBCN18 | Australia | 1988 | M1 | d; e; g; j; m |
| S7 | France | 2000 | HB-S7 | f; h; k; l; n |
| P27d | France | 1995 |  | l;o |
| Au4603 | Australia | 2015 |  | This study |
| Au7309 | Australia | 2015 |  | This study |
| D53 | Australia | 2016 |  | This study |
| 929 | Canada | 1993 | A929 | p; k |

<sup>a</sup> Kutcher et al. (2010); <sup>b</sup> Larkan et al 2016; <sup>c</sup> Ghanbarnia et al. 2018; <sup>d</sup> Balesdent et al 2002; <sup>e</sup> Balesdent et al 2005; <sup>f</sup> Zhang et al 2016; <sup>g</sup> Plissonneau et al 2017; <sup>h</sup> Petit Houdenot et al. 2019; <sup>i</sup> Balesdent et al. 2013; <sup>j</sup> Purwantara et al. 2000; <sup>k</sup> Leflon et al. 2007; <sup>l</sup> Somroo 2016; <sup>m</sup> Marcroft et al. 2012; <sup>n</sup> Brun et al. 2001; <sup>o</sup> Delourme et al. 2008; <sup>p</sup> Somda et al. 1999

**Table 3.**
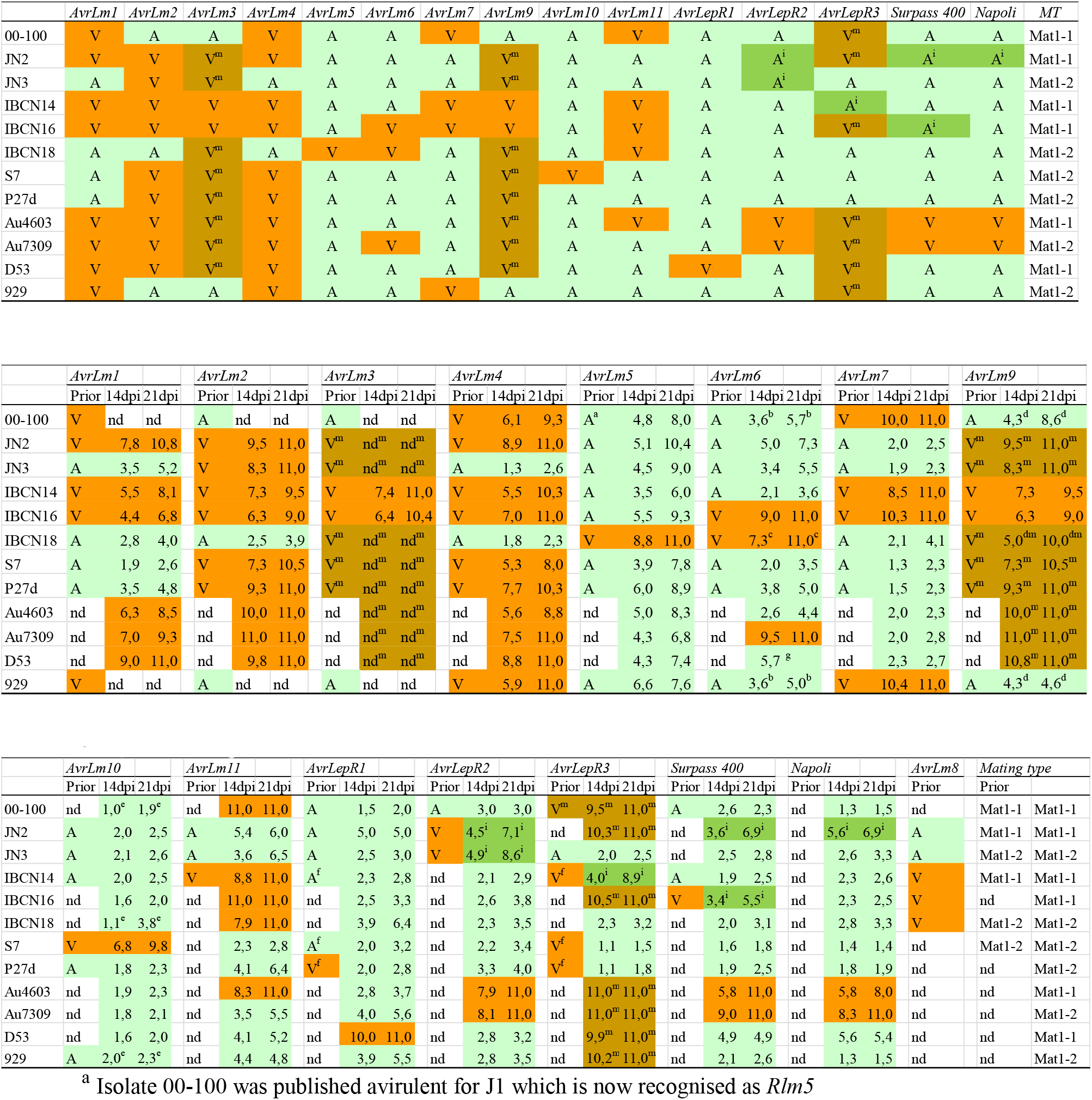
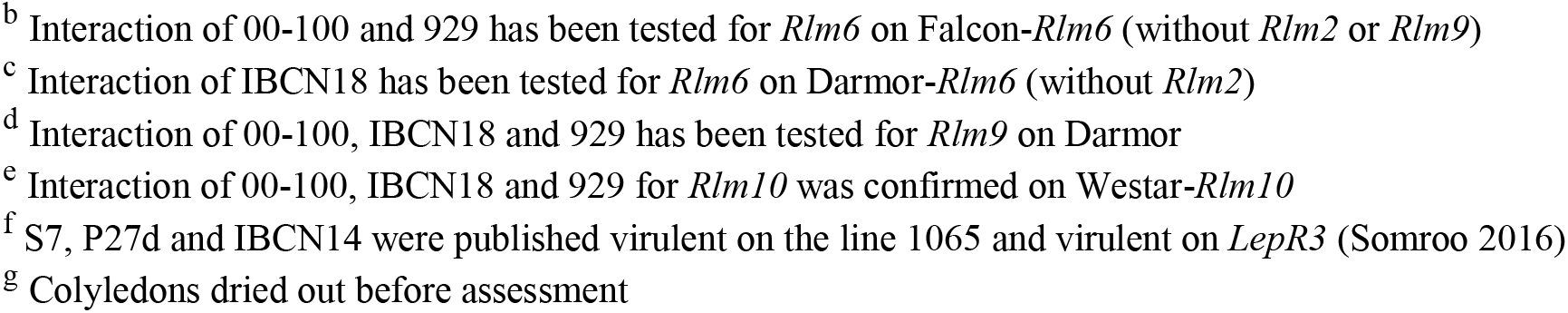
Arirulence profiles of the fungal reference isolates based on disease scores on seedlings of Brassica lines and varieties (listed in Table 1). **3A**. Summary of virulence (V), avirulence (A) and mating type (MT) alleles. **3B**. Detailed information and scores. Presence/absence of the avirulence allele in the Prior information is based on publications (Table 2) with “V” and orange (darker) highlight referring to the presence of the virulence allele; “A” and green (lighter) highlight referring to the presence of the avirulence allele; “nd” referring to alleles that were not determined; “V^m^” and brown highlight indicate that avirulence on Rlm7 masks the interaction with Rlm3 and Rlm9; and that virulence on Rlm1 confers virulence on LepR3; “Ai” with dark green highlight indicate intermediate (avirulent) interaction with larger symptoms than avirulent reference isolates, but smaller appearing later and darker than the virulent reference isolates. Scores at 14 dpi and 21dpi are means of 4 inoculation points x 6 plants. Mating type alleles were according to Cozjinsen & Howlett (2003).

### Scoring

Infection phenotypes were scored at 14 dpi and confirmed at 21 dpi as described in Chèvre et al. (2008). The level of cotyledon infection was scored on a 0 to 11 scale in which 0 = no disease and 11 = cotyledons collapsed from phoma stem canker. The scale was modified from the 0 to 9 scale described earlier in which 0 = no darkening around wound – typical response of non-inoculated controls or controls inoculated with water only; 1 = limited blackening around wound and lesion diameter of 0.5 to 1.5 mm; 3 = dark necrotic lesion with diameter of 1.5 to 3.0 mm; 5 = nonsporulating, 3 to 6 mm in diameter, lesion sharply delimited by darkened necrotic tissue; 7 = grayish green tissue collapse, lesion of 3 to 5 mm, with sharply delimited, non-darkening margin; and 9 = rapid tissue collapse at approximately 10 days, accompanied by profuse sporulation in large lesion (greater than 5 mm), and lesions with diffuse, non-darkened margins. To this scale an additional category of “11 = cotyledon totally collapsed, very often dry” was added (Chèvre et al., 2008) to categorize the most susceptible plants. Plants of classes 0 to 5 were considered as resistant as well as plants in classes 5 to 7 showing typical resistant plant reaction with blackening and limited symptoms that did not develop further between the two assessment dates at 14 and 21 days after inoculation. Plants of classes 9 to 11 were classified as susceptible, with symptoms appearing quickly, no blackening observed in lesions and with rapid disease progression between the two dates of assessment (14 and 21 days after inoculation). The scores were averaged over all infected plants.

## Results

### A complete set of isolates to characterize the presence / absence of 13 avirulence genes

With a set of 12 isolates, we were able to expand the previously available characterization on 13 of the 17 described resistance genes, having clearly compatible and clearly incompatible isolates for each interaction (Table 3A,B). There are only a few interactions that we were not able to confirm, and we thus rely on prior information (Table 3B). For isolates 00-100, 929 and IBCN18, avirulence allele AvrLm2 precluded conclusion on our differential lines for *Rlm3*. For isolates 00-100 and 929, avirulence allele AvrLm9 precluded conclusion on our differential lines for *Rlm1* and *Rlm2*. Conversely, the presence of the avirulence allele AvrLm9 in 00-100 together with virulence on Darmor*Rlm11* indicates the absence of *Rlm9* gene in Darmor *Rlm11*.

Expanding our set of isolates with clearly virulent ones allowed us to detect inconsistent behaviours or intermediate (avirulent) phenotypes. On the Topas-*LepR1* line, using isolate D53, we were able to obtain clearly compatible symptoms (greyish leaf spots, large and with an early increase in size), contrasting with the intermediate phenotype. We consider the intermediate phenotype as avirulent, because depending on the test, isolates may cause large lesions on some of the cotyledons but the lesions are dark and this intermediate phenotype is not consistent from one round of testing to the next. Discrepancies occurred for IBCN14, S7 and P27d published virulent on the 1065 line with *LepR1* (Somroo 2016) but having incompatible interactions on Topas-*LepR1* in our tests. On the Topas-*LepR2* line, using isolates Au4603 and Au7309, we were able to obtain clearly compatible symptoms (greyish leaf spots, large and with an early increase in size), contrasting with the intermediate phenotype of JN2 and JN3 (symptoms of large size, but dark as in incompatible interactions and with restricted increase in size). On the Topas-*LepR3* line, IBCN14 had intermediate phenotype. Discrepancies occurred for IBCN14, S7 and P27d published virulent on *LepR3* based on the use of Surpass 400 (Somroo 2016) but having incompatible interactions on Topas-*LepR3* in our tests. On the Surpass 400 and Napoli varieties, isolates Au4603 and Au7309 had clearly compatible symptoms (greyish leaf spots, large and with an early increase in size), whereas JN2 had intermediate phenotype on both varieties. IBCN14 had intermediate phenotype on Surpass 400.

### A current French population characterized for 13 avirulence gene frequencies

For the French *L. maculans* population sampled in 2018 at Le Rheu, we provide the current status for 13 avirulence frequencies (Table 4, Figure 1). Avirulence frequencies on the genes *Rlm1*, *Rlm2*, *Rlm3*, *Rlm4*, *Rlm7*, *Rlm9* and *LepR3* were low. In the opposite, avirulence were absent or very low for the genes *Rlm5*, *Rlm6*, *Rlm10*, *Rlm11*, *LepR1* and *LepR2* (Table 4, Figure 1). Noteworthy, in the population tested, most of the isolates avirulent on *Rlm7* had somewhat higher scores on Surpass 400 and Napoli than the ones virulent on *Rlm7* (Table 4).

**Table 4.**
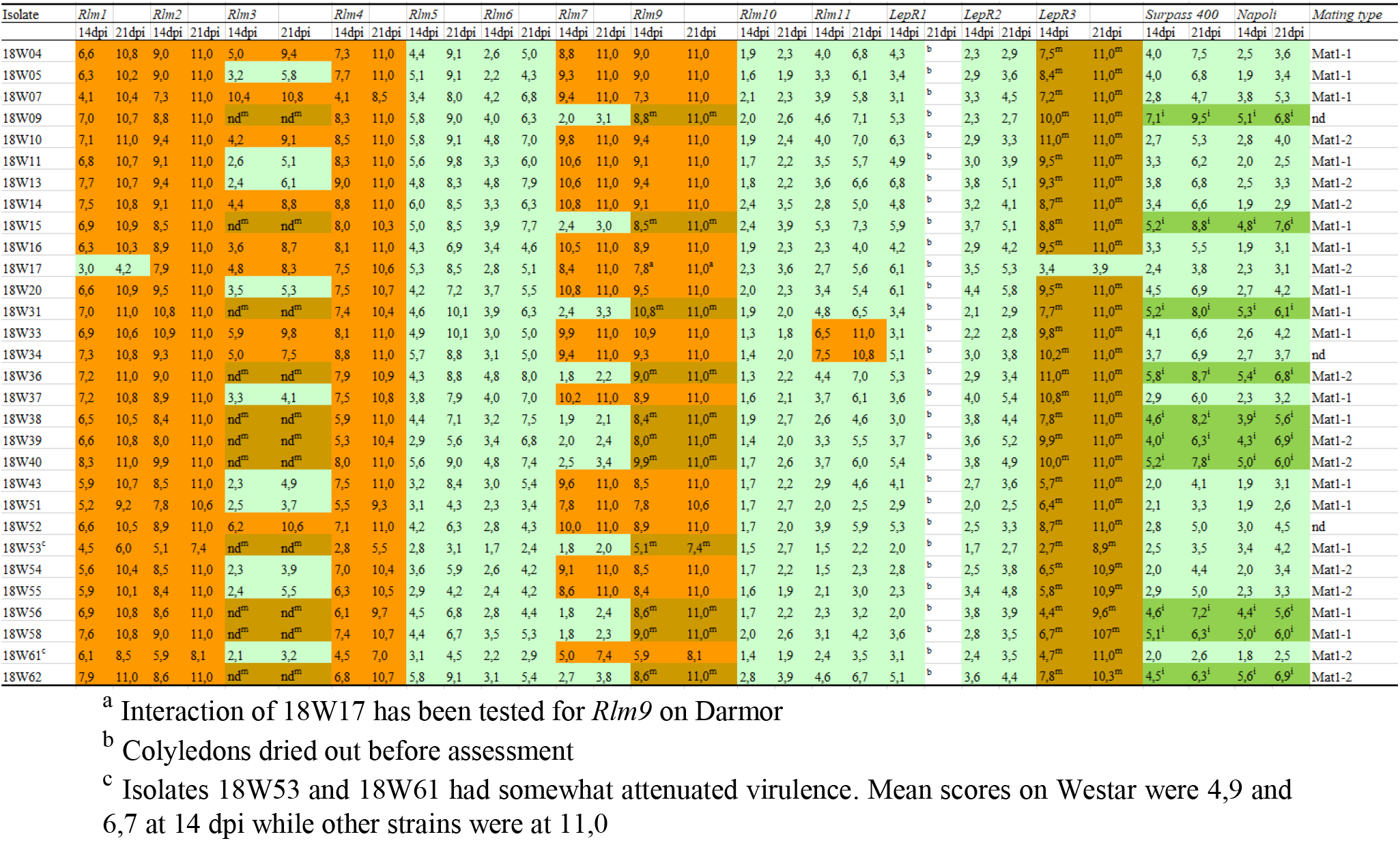
Presence/absence of the avirulence allele in the 2018 population sampled at Le Rheu based on disease scores on seedlings of Brassica lines and varieties (listed in Table 1) with orange (darker) highlight referring to the presence of the virulence allele; green (lighter) highlight referring to the presence of the avirulence allele; “nd” refering to alleles that were not determined; “^m^” and brown highlight indicate that avirulence on Rlm7 masks the interaction with Rlm3 and Rlm9; and that virulence on Rlm1 confers virulence on LepR3; “i” with dark green highlight indicate intermediate (avirulent) interaction with larger symptoms than avirulent reference isolates, but smaller appearing later and darker than the virulent reference isolates. Scores at 14 dpi and 21dpi are means of 4 inoculation points x 6 plants. Mating type alleles were according to Cozjinsen & Howlett (2003).

**Figure 1.**
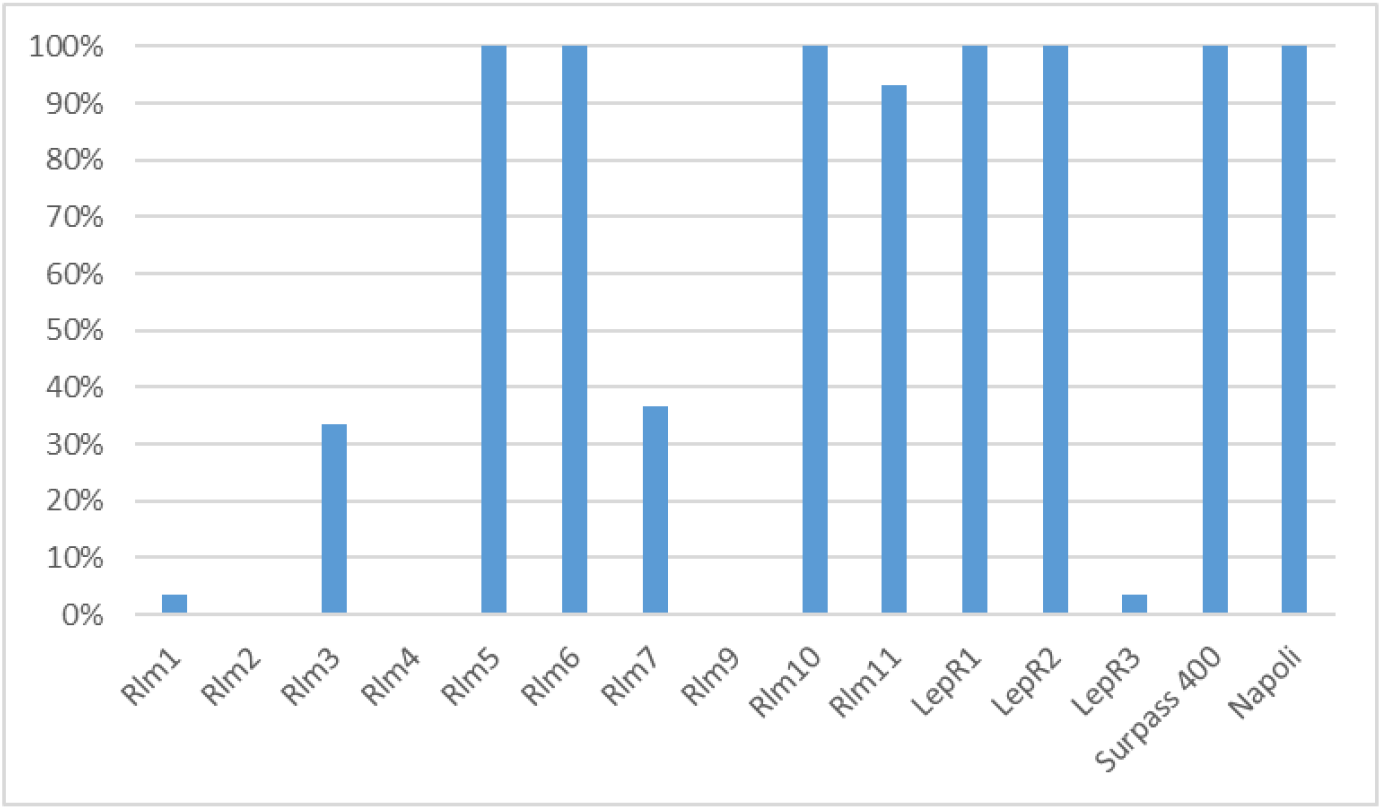
Avirulence frequencies on 13 resistance genes and two varieties, within a population of 30 isolates sampled at Le Rheu in 2018. See Table 1 for the differential lines used.

The two isolates 18W53 and 18W61 showed somewhat attenuated symptoms. Lesion size were smaller than other virulent isolates even on Westar without resistance genes (18W53 4.9 at 14 dpi and 10.1 at 21 dpi; 18W61 6.7 at 14 dpi and 11.0 at 21 dpi when other isolates were all at 10.0 or 11.0 at 14 dpi). Lesion appearance was sometimes darker, almost like avirulence symptoms. Such weakly virulent isolates were commonly found in the first generations of recurrent selection on very efficient resistant varieties in the field (Brun et al. 2000; 2010). It was quite unexpected to find some just sampling airborne *L. maculans* population, and their fitness could deserve further investigation.

## Discussion

### A complete set of isolates to characterize the presence / absence of 13 avirulence genes

With 12 isolates, we were able to expand the previously available characterization for 13 of the 17 described genes, having clearly compatible and clearly incompatible isolates for each interaction. A complete characterization of isolates was missing because different genes have been characterized by different teams, because new interactions are continuously identified (Larkan et al. 2019) and because seed of differentials are difficult to obtain. We choose isolates with already published information (Table 2) in order to increase the probability that they will already be available in different teams for further work. In most cases, our results were congruent with prior information (Table 3).

As opposed to prior information, none of the published isolates were clearly virulent on the lines Topas-*LepR1* and Topas-*LepR2* (Table 3). We thus expanded the set to encompass clear contrasted phenotypes. Isolate D53 was virulent on *LepR1* and isolates Au4603 and Au7309 were virulent on *LepR2*. Discrepancies with prior information might be due the use of different lines to test for *LepR1* and *LepR2*, because the genetic background of Topas (in lines Topas-*LepR1* and Topas-*LepR2*) confers higher resistance than the genetic background of Westar (in lines 1065 and 1135) (Larkan et al. 2016; Haddadi et al 2019). Also, on the Topas lines we repeatedly observed cotyledons drying out, even with incompatible interaction. This could have in some cases caused mis-scorings in previous studies.

When testing a population of isolates, it is necessary to include controls in each test, so that both compatible (virulent) and incompatible (avirulent) reactions are generated and the expression of interactions is confirmed. Isolates from this fully-tested set were used as controls in the second part of our study, with at least two reference isolates with known phenotypes per test.

### A current French population characterized for 13 avirulence gene frequencies

We provide the current status for 13 avirulence frequencies for the French *L. maculans* population sampled in 2018 at Le Rheu. Overall, as compared to previous studies, resistance genes deployed in varieties have lost efficacy and only avirulences corresponding to resistance genes not deployed retain efficacy. Previous large-scale surveys in France were undertaken in 2000-2001 with phenotyping for AvrLm1 to AvrLm9 (Balesdent et al. 2006), later completed with genotyping for AvrLm11 (Balesdent et al. 2013) and specific studies on AvrLm3 and AvrLm7 (Leflon 2013; Balesdent et al. 2015; 2019). Further information on the local population at Le Rheu is available in 2004-2005 with phenotyping for AvrLm1; AvrLm4; AvrLm5 and AvrLm6 (Brun et al. 2010). Based on the high corresponding avirulence frequencies, the genes *Rlm1*, *Rlm2*, *Rlm3*, *Rlm4*, *Rlm7*, *Rlm9* and *LepR3* are not expected to provide sufficient resistance if used in varieties. This indicates that so far all the resistance genes used in France on significant acreage without additional management practices have lost efficacy. *Rlm2*, *Rlm4* and *Rlm9* being present in former varieties (Rouxel et al. 2003a) have lost efficacy against French populations long ago (Rouxel et al. 2003b), followed by *Rlm1* and *Rlm7* deployed later (Rouxel et al. 2003b; Balesdent et al. 2015; 2019; Leflon 2013). Whereas in Canada, *Rlm3* has lost efficacy following its use in varieties (Liban et al. 2016; Zhang et al. 2016), this interaction was masked in France at a time when most isolates were avirulent on *Rlm7*, then regained a slight efficacy but isolates virulent on both *Rlm3* and *Rlm7* are now increasing in frequency (Leflon 2013; Balesdent et al. 2015; 2019).

Based on the low corresponding avirulence frequencies, the genes *Rlm5*, *Rlm6*, *Rlm10*, *Rlm11*, *LepR1* and *LepR2* are expected to filter out most of *L. maculans* isolates, thus lowering disease levels on crops. An optimistic point of view would conclude that there are ample resources for breeding oilseed rape varieties resistant to French *L. maculans* populations. However, first these results need confirmation on additional populations, and second, documented cases from other countries indicate that these genes are likely to be overcome if deployed. In Australia, isolates virulent on *Rlm5* increased in frequency following the used of juncea canola (Elliott et al. 2015; Van de Wouw et al. 2017). Further, isolates virulent on *Rlm6* have increased in frequency even before the deployment of the corresponding resistance gene (Van de Wouw et al. 2017). In France, *Rlm6* was overcome within a few years in field experiments (Brun et al. 2010). The specific case of Surpass 400 variety and *LepR1*, *LepR2* and *LepR3* genes is discussed in the following section. What remains on the list? *Rlm10* and *Rlm11*, however virulent isolates do exist in French populations (Somda et al. 1999; Balesdent et al. 2013). The *Rlm10* gene was introduced from *B. nigra* and was not deployed so far in oilseed rape varieties. Avirulence for AvrLm10 was seldom tested on large scale, so information is scarce. Two alleles contribute to the interaction (Petit-Houdenot et al. 2019). Avirulence frequency was high in French field experiments even though virulent isolates did exist in populations (Brun et al. 2001). All Canadian isolates tested were avirulent (Kutcher et al. 2010). In Australia, *B. nigra* lines retained efficacy in 2004-2005 (Light et al. 2011). Avirulence for AvrLm11 was tested at population scale only in France (Balesdent et al. 2013). Isolates lacking the minichromosome and the AvrLm11 markers were present at low frequencies in 2000 (4,5%) and remained so in 2010 (2,4%). Avirulence was high (93%) in our study, congruent with the – as far as we know – absence of the gene in oilseed rape varieties. Recent results document the two genes *LepR5* and *LepR6* (Larkan et al. 2019), but their efficacy and potential interactions or allelism with other genes are not yet documented.

### *Comments on the Surpass 400 variety,* RlmS, LepR2 *and* LepR3 *genes discussion, and discrepancies for* LepR1 *and* LepR2

All isolates at le Rheu were avirulent on the Topas-*LepR1* and Topas-*LepR2* lines, and all but one were virulent on the Topas-*LepR3* (this isolate being avirulent on *Rlm1*). Comparisons with prior information is difficult because of two reasons.

On the one hand, the variety Surpass 400 was deployed in Australia and has lost efficacy overtime (Li et al. 2003). Concomitant decrease in AvrLm1 was observed (van de Wouw et al. 2017). Over years, there was agreement on the presence of two resistance genes in Surpass 400, but much debate about the genes being *LepR2* and *LepR3* (Yu et al. 2008), *Rlm1* and *RlmS* (Marcroft et al. 2012), *Rlm1* and *LepR3* (Van de Wouw et al. 2017), *RlmS* and *LepR3* (Liban et al. 2016). The consequence is that when Surpass 400 was used as a differential, inferences depended on the assumptions on the genes present. To our knowledge, the debate is currently settling, as discussed at workshops in recent international conferences. However, the confirmation that *RlmS* is *LepR2* still awaits publication, that our study is definitely not designed to do. Following this hypothesis, one of the genes in Surpass 400 would thus be *LepR2*. Similarly, our study is not sufficient to distinguish between either *Rlm1* or *LepR3* being present in Surpass 400. However, phenotypically, isolates virulent on *Rlm1* would be compatible also on *LepR3* (Larkan et al. 2013). This is the reason why we come back to the idea that Surpass 400 has *LepR2* and *LepR3* (Table 1; Yu et al. 2008). In our tests, results are congruent with the presence of *LepR2* and *LepR3* in Surpass 400 (see isolates Au4603 and Au7309; Table 3).

On the other hand, the confusion was also amplified by the use of different lines to test for *LepR1* and *LepR2*. The genetic background of Topas (in lines Topas-*LepR1* and Topas-*LepR2*) confers higher resistance than the genetic background of Westar (in lines 1065 and 1135) (Larkan et al. 2016; Haddadi et al 2019). Also, on the Topas lines we repeatedly observed cotyledons drying out, even with incompatible interaction. This could have in some cases caused mis-scorings. These two causes might explain the discrepancies between prior information and our results on reference isolates (Table 3).

Based on these very low corresponding avirulence frequencies at Le Rheu, the genes *LepR1* and *LepR2* are expected to filter out most of *L. maculans* isolates, thus lowering disease levels on crops. In France, to our knowledge, the *LepR1* gene is not present in varieties so far, and there was no prior knowledge about French populations. Varieties with *RlmS*, such as Napoli, are now becoming available in France (Terres-Innovia 2019). *RlmS* is postulated to be *LepR2* (discussed above) and our results concerning *LepR2* and Napoli are congruent. Because the strains Au4603 and Au7309 virulent on LepR2 are avirulent on *Rlm5*, *Rlm6*, *Rlm7*, *Rlm10*, *Rlm11*, *LepR1* we can confirm the absence of these genes in Napoli. Further investigations are needed to ascertain the absence of the other *Rlm* genes.

Despite the current very low avirulence frequencies on *LepR1* and *LepR2*, documented cases from other countries indicate that these genes are likely to be overcome if deployed without additional management practices. In Australia, Surpass 400 (likely containing *LepR2* and *LepR3,*) has lost efficacy over time (Li et al. 2003; Sprague et al. 2006; Van de Wouw et al. 2017). Field experiments in Australia indicates that isolates virulent on *LepR1* are present in populations and the resistance gene does filter out avirulence isolates, thus imposing selection pressure on pathogen populations (Bousset et al. 2018). In Canada, for *LepR1* avirulence frequency was 15% in 2010 (Liban et al. 2016; line 1035); 40% in 2012 (Zhang et al. 2016; line 1065); from 10 to 85% in 2012-2014 depending on field and region (Somroo et al. 2016; line 1065). For *LepR2*, avirulence frequency was absent in 2010 (Liban et al. 2016; line 1065); 12% in 2012 (Zhang et al. 2016; line 1135); from 0 to 20% in 2012-2014 depending on field and region (Somroo et al. 2016; line 1135). For Surpass 400 (postulated to have *LepR3* and *RlmS* in these studies), avirulence frequency was around 50% in 2010 (Liban et al. 2016); 35% in 2012 (Zhang et al. 2016) from 0 to 90% in 2012-2014 depending on field and region (Somroo et al. 2016). Avirulence on *LepR3* was 98% in Canada (Kutcher et al. 2010); however, it was postulated from tests on Surpass 400 considered having *Rlm1* and *LepR3*. Thus this has to be reinterpreted as frequencies of the virulence on *LepR2* plus *LepR3* combination. Taken together, these informations call for care in the deployment of the currently efficient resistance genes.

### Options for a more durable use of resistance genes

While the call to wisely manage the available host resistance genes is not recent, it is still relevant. Importantly, the loss of efficacy of disease resistance genes can be averted when recommendation for deployment are produced on time (Van de Wouw et al. 2014). When local increase in disease severity on sentinel plots was detected for the variety Hyola 50 in the Eyre Peninsula, Australia, recommendations of not sowing the varieties at risk in this area were followed, preserving their efficacy countrywide.

Without additional management strategies, the wider the acreage of the variety is, the faster and larger the increase in frequency of the corresponding virulence could be (Bousset & Chèvre, 2013). The deployment of different resistance genes through space and time drives the genetic composition of the pathogen population (Hovmøller et al., 1997; Papaïx et al., 2011). Modelling studies (Lô-Pelzer et al. 2010) and field experiments provide knowledge on the interplay between the acreages on which genes are deployed, the combination of genes in varieties, the intensity of inoculum sources and the spatial organization of crops in the landscape. There is currently no agreement on the best way to deploy available resistance genes, either individually or stacked in varieties (Rimbaud et al. 2018). On the one hand, diversifying selection has been proposed to play on pathogen’s weakness, such as stabilizing evolutionary dynamics (Zhan et al., 2015). On the contrary, stacking genes or QTLs in pyramids is also advocated (Pilet-Nayel et al. 2017). It is worth noticing that while stacking might be of interest when the corresponding virulences are absent from pathogen populations (Lof et al., 2017), this strategy leaves no possibility of later leveraging the decrease of unnecessary virulences. Based on theoretical grounds in evolutionary biology of plant pathogen coevolutionary dynamics, it has been proposed that selectively reducing the contribution of pathogen populations from fields cropped with a resistant variety to the initial inoculum of the following season could slow down adaptation (Bousset & Chèvre, 2013). Field experiments with contrasting amounts of inoculum and contrasting levels of pre-adaptation (simulated by mixing stubble sources) confirm this hypothesis (Bousset et al., 2018). Thus, stabilizing the evolutionary dynamics should be considered, using options for reducing spore sources and reducing transmission by the spatial organization of crops.

Three kinds of management practices can be added to the deployment of resistance genes in order to reduce inoculum carry-over from one growing season to the next and lower population sizes. On the one hand, *L. maculans* survives on stubble. In field experiments, the reduction of inoculum by stubble management delayed adaptation (Daverdin et al. 2012). Currently, the previous year’s stubble is the primary contributing source for *L. maculans* (Marcroft et al., 2004) although changes in tillage cropping practice (McCredden et al., 2017) or shorter rotations might alter this situation. Options for reducing inoculum sources could include stubble management by burial (Marcroft et al., 2004), flooding or chemical application. Where possible, applying stubble management to reduce disease pressure on the following crops should be considered. On the other hand, wind dispersed ascospore produced by sexual reproduction are the main source of inoculum for *L. maculans* epidemics. Landscape structure is important for *L. maculans* transmission, such that increased isolation of crops (up to 500m) from any canola crops grown in the preceding year was associated with lower levels of disease (Marcroft et al., 2004). Transmission between fields can be predicted from spore dispersal (Marcroft et al., 2004; Bousset et al., 2015). Thus, spatially explicit models can be used to study and ultimately design combinations of landscapes, varietal choice and tillage practices promoting resistance durability against phoma stem canker (Lô-Pelzer et al., 2010; Rimbaud et al. 2018). As to the practicalities of spatial organization of the landscape, lessons have to be learned from the available case studies. Finally, combining qualitative and quantitative resistance within host lines has been shown to delay the loss of efficacy of qualitative resistance (Brun et al. 2010; Delourme et al. 2014). Because erosion of quantitative resistance has not been observed so far, deploying resistance genes in partially resistant backgrounds is a minimal precaution.

## Acknowledgements

**Acknowledgements**

We thank Susan Sprague for providing the *L. maculans* Au4603 and Au7309 isolates and Randy Kutcher for providing the *L. maculans* 00-100 isolate. We thank Martin Frauen from NPZ for providing the Napoli variety. We thank Gary Peng and Hossein Bohran for providing the Topas-*LepR1*; Topas-*LepR2 and* Topas-*LepR3* near isogenic lines. We thank Lucie Mieuzet for technical assistance. This work benefited from the financial support of INRA – the French National Institute for Agronomical Research. The authors declare the absence of conflict of interest. ME carried out experiments, LB and RD conceived and designed the study and prepared the manuscript, read and approved by all authors.

